# Single nucleotide polymorphisms associated with cytotoxicity of *Bacillus cereus* group strains in Caco-2 cells

**DOI:** 10.1101/2025.02.14.638278

**Authors:** Cassidy Prince, Taejung Chung, Kayla Kimble, Tyler Chandross-Cohen, Martin Wiedmann, Sophia Johler, Jasna Kovac

**Affiliations:** Department of Food Science, The Pennsylvania State University, University Park, 16802 PA, United States; Department of Food Science, Cornell University, Ithaca, 14853 NY, United States; Institute for Food Safety and Hygiene, Vetsuisse Faculty, University of Zurich, 8057 Zurich, Switzerland

**Keywords:** *Bacillus cereus*, cytotoxicity, Caco-2, whole-genome sequencing, enterotoxin, Nhe, Hbl, CytK

## Abstract

*Bacillus cereus sensu lato* (*s.l*.) encompasses strains with diverse impacts, ranging from foodborne illness and anthrax to beneficial applications in agriculture and industry. While the risk of anthrax and emetic intoxication can be reliably predicted by the presence of specific virulence genes, predicting diarrheal foodborne illness risk based solely on enterotoxin gene presence has proven unreliable. In this study, we evaluated cytotoxicity against Caco-2 human gut cells using a diverse collection of *B. cereus s.l.* isolates representing all eight *panC* phylogenetic groups and conducted genomic analyses to identify predictive markers of cytotoxicity. Isolates from *panC* groups I, IV, and V exhibited significantly higher cytotoxicity compared to other groups, although individual isolates from other *panC* groups have also been linked to illness. Logistic and random forest regression models revealed that while the presence of enterotoxin genes was a sensitive indicator of cytotoxicity, it lacked specificity. Logistic regression analysis identified 21 nonsynonymous single nucleotide polymorphisms (SNPs) within enterotoxin (Nhe and Hbl) gene sequences that were more effective predictors of cytotoxicity, providing higher specificity with comparable sensitivity. These SNPs achieved accuracy and precision values exceeding 0.7. Random forest models highlighted the importance of *panC* group, enterotoxin gene SNPs, and the presence of the full *hbl* operon as key predictors of cytotoxicity. The strong sensitivity, specificity, and biological relevance of these SNPs position them as promising markers for improving strain-based risk assessment of *B. cereus s.l*.

**IMPORTANCE:** Enterotoxin genes have been associated with *B. cereus sensu lato* (*s.l.*) diarrheal foodborne illness; however, their mere presence in a genome is an unreliable predictor of an isolate’s cytotoxicity towards human gut epithelial cells. To improve food safety risk assessment, more specific markers of cytotoxicity are required. In this study, we identified nonsynonymous SNPs within the coding sequences of the enterotoxins Nhe and Hbl. These SNPs offer potential targets for rapid molecular tests to identify *B. cereus s.l.* isolates with an elevated food safety risk due to their capacity to inflict cytotoxic damage on human gut epithelial cells. Implementation of such markers upon validation could improve consumer safety while reducing food waste.

## INTRODUCTION

The *Bacillus cereus* group, or *Bacillus cereus sensu lato* (*s.l.*), comprises multiple genomospecies of spore-forming bacteria that are widespread in nature. *B. cereus s.l.* includes strains with diverse impacts, ranging from clinically significant pathogens to beneficial bioinsecticides. Pathogenic strains are associated with emetic (1) and diarrheal foodborne illnesses (2, 3), fatal anthrax and anthrax-like disease (4, 5), meningitis (6), and skin infections (2). Conversely, beneficial strains are used commercially for their production of insecticidal crystal proteins (7, 8). Accurate differentiation between harmful and beneficial strains is crucial for public health.

Existing PCR- or genome-based detection methods effectively identify strains linked to emetic toxin (cereulide)-mediated intoxication, based on the presence of the *ces* gene cluster, or anthrax-like disease, based on anthrax toxin-encoding genes (*cya, lef, pagA*) (9). However, predicting diarrheal foodborne toxicoinfection remains challenging, as all *B. cereus s.l.* isolates harbor at least one enterotoxin gene, and diarrheal illness has been linked to isolates with varying enterotoxin gene profiles (2, 10–12).

The primary enterotoxins implicated in diarrheal illness are nonhemolytic enterotoxin (Nhe), hemolysin BL (Hbl), and cytotoxin K-1/K-2 (CytK-1, CytK-2). Nhe and Hbl are tri-partite pore-forming toxins encoded by *nheC*, *nheB*, *nheA*, and *hblA*, *hblD*, and *hblC*, respectively, with HblA initiating pore formation with NheC initiating pore formation (13, 14, 15). Both toxins lyse host cells by inducing potassium efflux and activating the NLRP3 inflammasome (16, 17). CytK, a β-barrel-forming toxin encoded by *cytK-1* or *cytK-2*, is associated with cell lysis, though its mechanism remains unclear (18, 19). CytK-1 is exclusive to *B. cytotoxicus*, while CytK-2 is found across multiple *B. cereus s.l.* species (10, 11). Additional virulence factors, including sphingomyelinase (SMase), phosphatidylinositol-specific phospholipase C (PI-PLC), immune inhibitor A1 (InhA1), and neutral protease (NprA or NprB) may act synergistically with enterotoxins to enhance cytotoxicity (20–25).

Despite the universal presence of enterotoxin genes, not all *B. cereus s.l*. isolates exhibit cytotoxicity toward human cell lines (10, 11). Cytotoxicity varies among genomospecies and phylogenetic groups without aligning strictly to any specific clade (10, 11). This unpredictability underscores the need for improved genetic markers to identify high-risk isolates capable of causing toxicoinfection.

Previous studies have primarily investigated cytotoxicity in relation to *panC* phylogenetic groups or the presence of known virulence genes (10, 26, 27). However, the role of sequence variations within enterotoxin genes remains unexplored. To address this gap, we determined and analyzed the cytotoxicity of 270 phylogenetically diverse *B. cereus s.l.* isolates from all eight *panC* phylogenetic groups on Caco-2 cells. Using logistic and random forest regression analyses, we identified 21 nonsynonymous SNPs within enterotoxin sequences that were strongly associated with increased cytotoxicity. These SNPs were further evaluated for diagnostic metrics, including accuracy, precision, sensitivity, specificity, and permutation variable importance. Unlike conventional PCR-based gene detection, our whole-genome sequencing approach allowed for the identification of functionally significant SNPs that may directly influence cytotoxicity, paving the way for more precise diagnostic tools in food safety.

## RESULTS & DISCUSSION

### Diverse *B. cereus s.l.* isolates exhibit a broad range of cytotoxicity toward Caco-2 cells

We analyzed 270 phylogenetically diverse *B. cereus s.l.* isolates (Fig. 1) from various sources, including food (n=224), soils or sediments (n=36), commercial biopesticides (n=3), clinical samples (n=2), and unknown sources (n=5). These isolates were previously identified in multiple studies (3, 11, 28–32), with 240 isolates whole-genome sequenced for the first time in this study. To ensure diversity and avoid clonality, all isolates included in the study differ by more than 5 core genome SNPs. Phylogenetic grouping and genomospecies assignment were performed using BTyper3 (9, 33). All eight *panC* phylogenetic groups were represented (Fig. 1, Table 1), including one isolate identified as *Bacillus clarus*, a newly characterized effective species (34). Metadata and NCBI SRA accession numbers are listed in the Supplemental Material Table S1.

**FIG 1.**
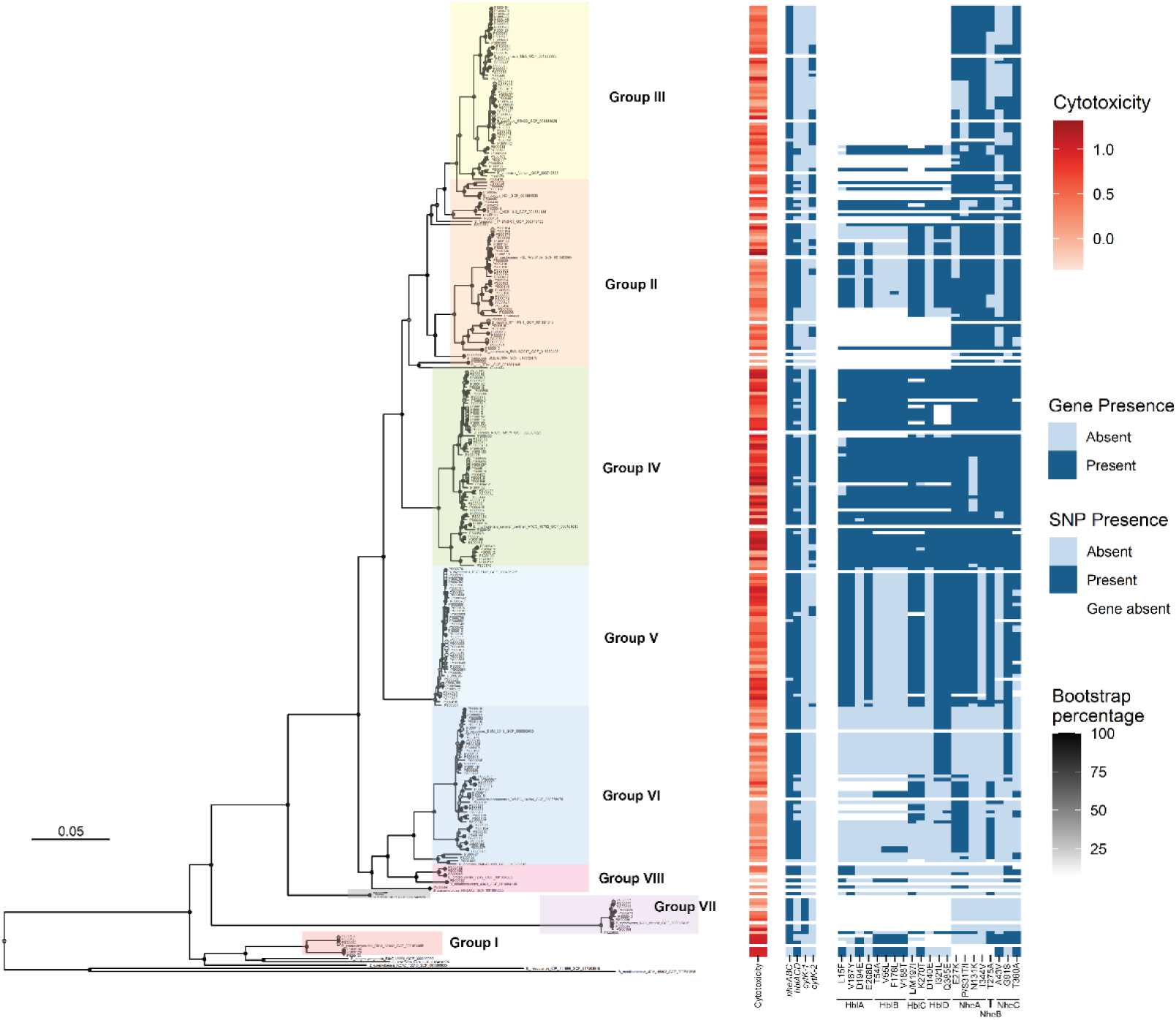
Maximum-likelihood phylogenetic tree of the 270 studied isolates and type strains with cytotoxicity values and enterotoxin gene and SNP presence. The tree was constructed using core genome SNPs identified with kSNP v2.13. *panC* phylogenetic groups are highlighted in different colors, and the *B. clarus* isolate is highlighted in gray. Type strain genomes are labelled with the corresponding species name and have exclusively white bars in the heatmap. Bootstrap percentages are displayed as colored circles at nodes, with darker circles indicating higher bootstrap percentages. From left to right, the heatmaps denote cytotoxicity, presence of a full enterotoxin gene operon, and presence of the cytotoxic SNP variant. White bars in the SNP heatmap indicate that the gene in which the SNP was detected is absent. Dark blue bars in the SNP heatmap indicate the presence of the amino acid associated with higher cytotoxicity, and light blue bars indicate the presence of an amino acid associated with a lower cytotoxicity.

**TABLE 1.**
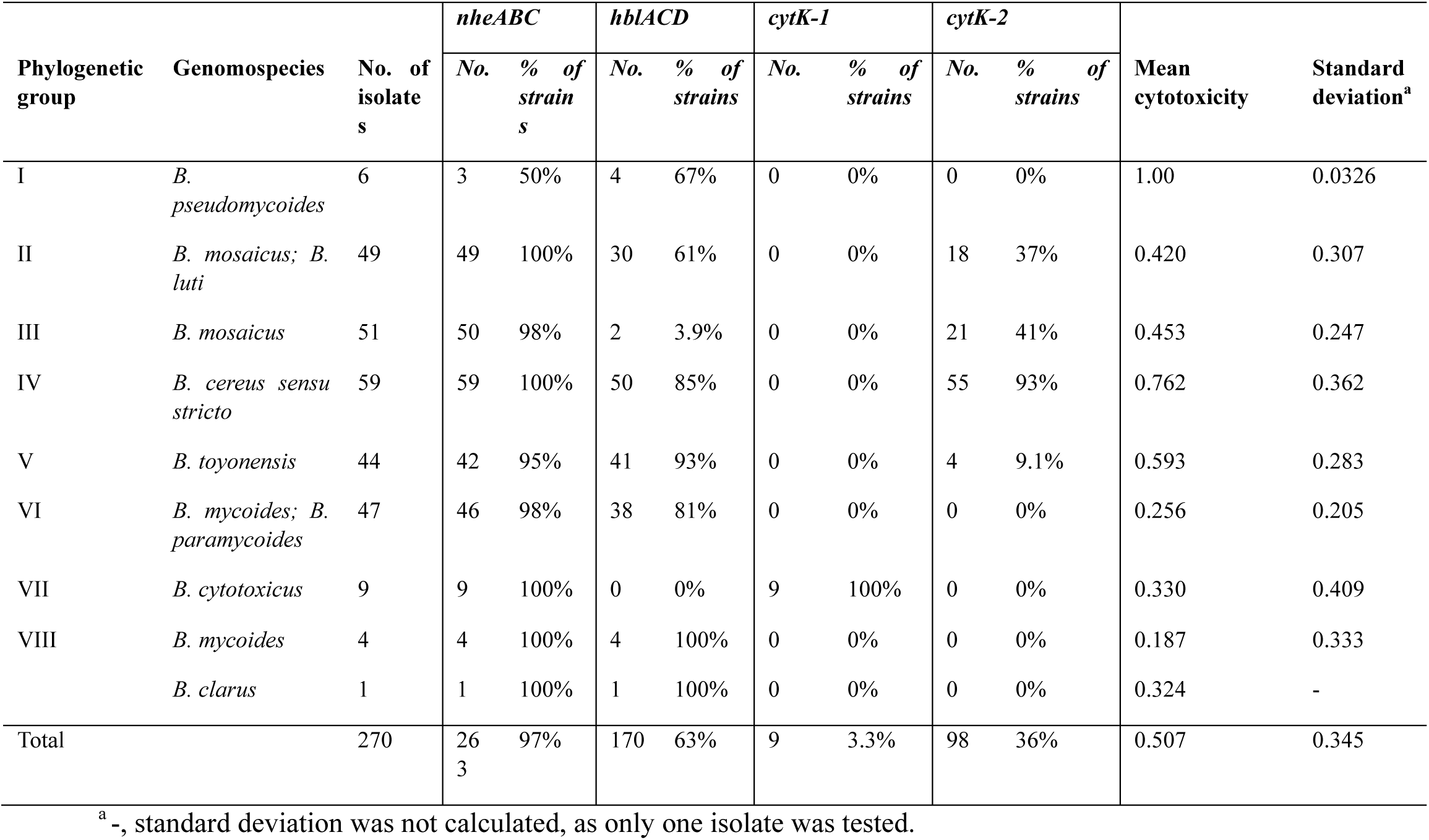
Mean cytotoxicity and enterotoxin gene presence by *B. cereus s.l.* phylogenetic group. Genes were detected with BTyper3 using 70% query coverage and 50% nucleotide sequence identity. Operons were considered present if all listed genes were detected.

| Phylogenetic group | Genomospecies | No. of isolates | *nheABC* |  | *hblACD* |  | *cytK-1* |  | *cytK-2* |  | Mean cytotoxicity | Standard deviation<sup>a</sup> |
| --- | --- | --- | --- | --- | --- | --- | --- | --- | --- | --- | --- | --- |
|  |  |  | No. | % of strains | No. | % of strains | No. | % of strains | No. | % of strains |  |  |
| I | *B. pseudomycoides* | 6 | 3 | 50% | 4 | 67% | 0 | 0% | 0 | 0% | 1.00 | 0.0326 |
| II | *B. mosaicus*; *B. luti* | 49 | 49 | 100% | 30 | 61% | 0 | 0% | 18 | 37% | 0.420 | 0.307 |
| III | *B. mosaicus* | 51 | 50 | 98% | 2 | 3.9% | 0 | 0% | 21 | 41% | 0.453 | 0.247 |
| IV | *B. cereus sensu stricto* | 59 | 59 | 100% | 50 | 85% | 0 | 0% | 55 | 93% | 0.762 | 0.362 |
| V | *B. toyonensis* | 44 | 42 | 95% | 41 | 93% | 0 | 0% | 4 | 9.1% | 0.593 | 0.283 |
| VI | *B. mycoides*; *B. paramycoides* | 47 | 46 | 98% | 38 | 81% | 0 | 0% | 0 | 0% | 0.256 | 0.205 |
| VII | *B. cytotoxicus* | 9 | 9 | 100% | 0 | 0% | 9 | 100% | 0 | 0% | 0.330 | 0.409 |
| VIII | *B. mycoides* | 4 | 4 | 100% | 4 | 100% | 0 | 0% | 0 | 0% | 0.187 | 0.333 |
|  | *B. clarus* | 1 | 1 | 100% | 1 | 100% | 0 | 0% | 0 | 0% | 0.324 | - |
| Total |  | 270 | 263 | 97% | 170 | 63% | 9 | 3.3% | 98 | 36% | 0.507 | 0.345 |

<sup>a</sup> -, standard deviation was not calculated, as only one isolate was tested.

Genes encoding Nhe, Hbl, CytK-1, and CytK-2 enterotoxins were identified using BTyper3 (33). All isolates’ genome assemblies contained at least one *nhe* gene, while only 73% contained at least one *hbl* gene. The full *nheABC* and *hblACD* operons were present in 97% and 63% of genomes, respectively (Table 1). The *hblACD* operon was largely absent from *panC* groups III and VII. *cytK-1* was exclusive to group VII genomes, while *cytK-2* was prevalent in groups II (37% of Group II isolates), III (41%), and IV (93%), but rare in group V (9.1%) and not detected in groups I, VI, VII, and VIII (Table 1). These findings align with existing literature (10, 12, 35). Multiple enterotoxin operons or genes were found in 212 genomes (79%). Phylogenetic groupings generally correlated with enterotoxin gene presence, indicating that phylogeny and enterotoxin gene presence covary (Table 1, Fig. 1).

Enterotoxin protein sequences showed variable conservation. CytK-1 and CytK-2 were highly conserved, with amino acid sequence identities of 99.7% and 86.3%, respectively. In contrast, other proteins showed lower sequence identities: NheB (72.5%), HblD (67.2%), NheA (57.8%), HblA (61.1%), HblB (54.4%), HblC (53.1%), and NheC (50.7%).

All isolates were tested for cytotoxicity using cell-free supernatants from early stationary-phase cultures, a time point when diarrheal enterotoxin secretion peaks (36). Caco-2 cells were exposed to the supernatants, and cytotoxicity was assessed using a WST-1 colorimetric assay (37). Absorbance values were normalized within plates using min-max normalization (38), with 0 representing sterile brain heart infusion (BHI) medium and 1 representing the cytotoxicity of the supernatant of the *B. cereus s.s.* ATCC 14579 type strain. High-cytotoxicity isolates caused visible cell lysis and detachment, while low-cytotoxicity isolates had minimal impact on cells (Fig. 2A).

**FIG 2.**
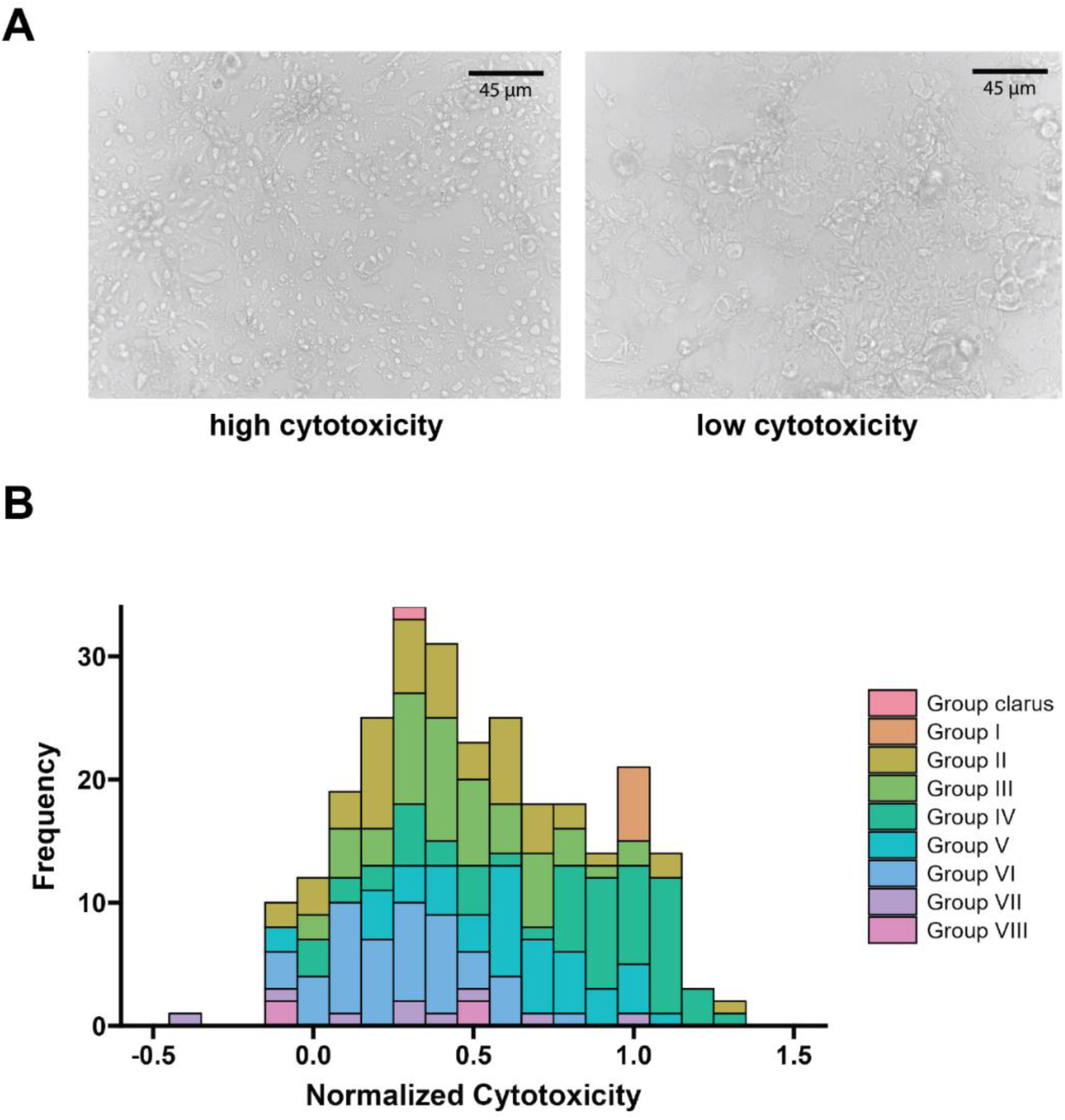
Cytotoxicity distribution of 270 isolates, as determined by intoxication of Caco-2 cells using cell-free supernatants. A. Microscope images of Caco-2 cells exposed to highly cytotoxic (normalized cytotoxicity of 1) or not cytotoxic (normalized cytotoxicity of 0) *B. cereus s.l.* supernatants. **B.** Histogram of normalized cytotoxicity values colored by *panC* group. A normalized cytotoxicity value of 0 indicates cytotoxicity equal to the noncytotoxic BHI control, and a value of 1 indicates cytotoxicity equal to the cytotoxic *B. cereus s.s.* ATCC 14579 supernatant control.

The mean normalized cytotoxicity across all isolates was 0.51 ± 0.35, indicating a metabolic activity reduction to approximately half that of ATCC 14579 strain. Some isolates exceeded a cytotoxicity value of 1, suggesting greater cytotoxicity compared to ATCC 14579, while others exhibited values below 0, suggesting enhanced metabolic stimulation compared to sterile BHI. The distribution of cytotoxicity values (Fig. 2B) highlights substantial variability among isolates, indicating differences in virulence potential within *B. cereus s.l.* A Shapiro-Wilk normality test confirmed that the cytotoxicity values were not normally distributed (Fig. 2B). This non-normal distribution suggests the possibility of defining a cytotoxicity threshold to distinguish “cytotoxic” from “non-cytotoxic” isolates. However, additional data, particularly from underrepresented phylogenetic groups, may influence the observed distribution and bring it closer to normality.

### Isolates from *panC* groups I, IV, and V exhibit significantly higher cytotoxicity to Caco-2

Previous studies have reported variation in cytotoxicity among *panC* phylogenetic groups (10, 11). To validate these findings, we compared cytotoxicity values across our collection of isolates. Group VIII isolates exhibited the lowest average cytotoxicity (0.187 ± 0.333), while group I isolates showed the highest (1.00 ± 0.0326) (Table 1, Fig. 3). Most *panC* groups, except group I, demonstrated broad distributions in cytotoxicity, as indicated by the large standard deviations (Table 1, Fig. 3).

**FIG 3.**
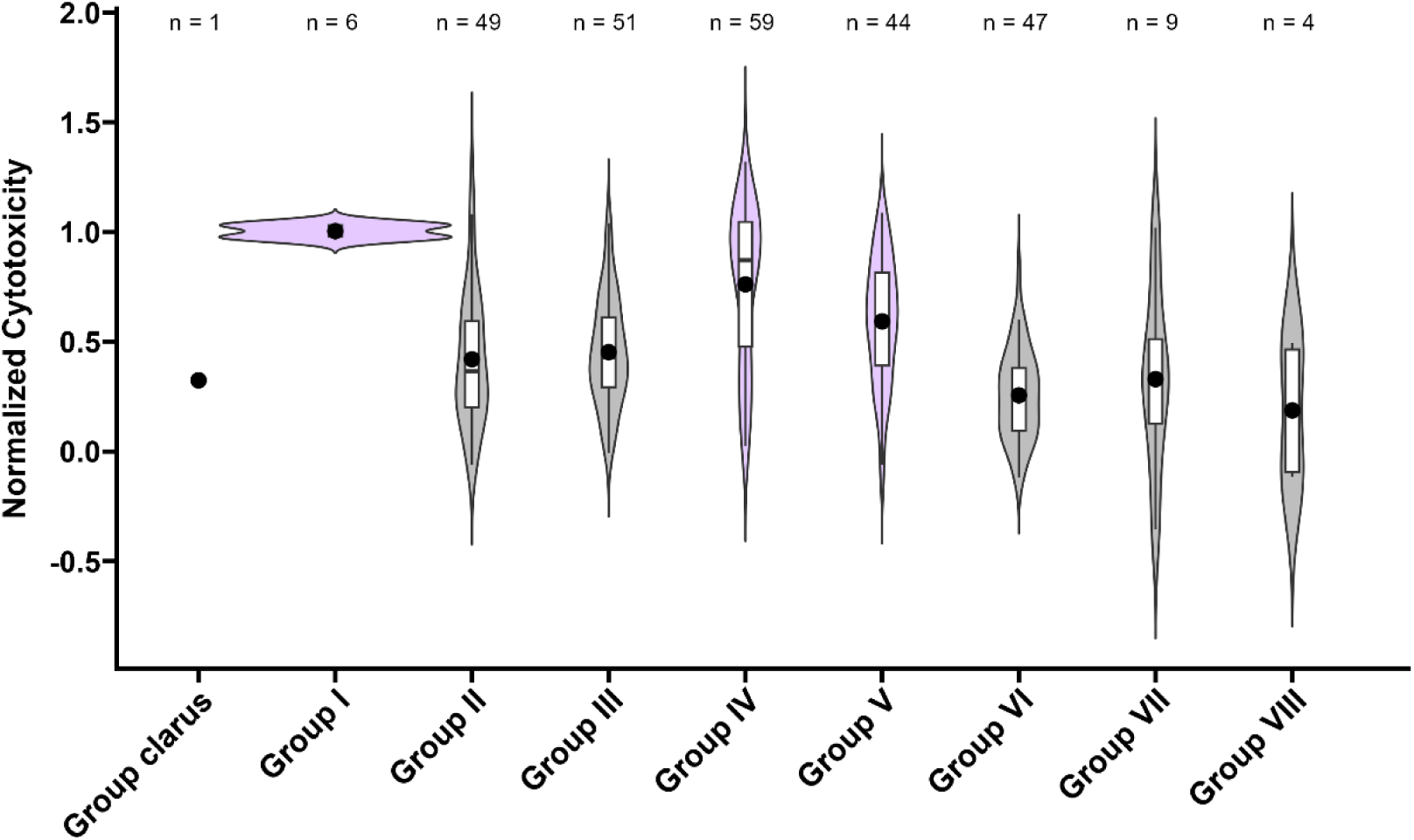
Cytotoxicity of isolates within individual phylogenetic groups. The black dot indicates the mean cytotoxicity per group. Groups I, IV, and V are colored pink, denoting significantly higher cytotoxicity compared to several other groups. The numbers of isolates tested per group are listed above each plot.

Since specific enterotoxin genes, such as *cytK-1*, are disproportionately represented in certain *panC* groups, we investigated whether *panC* group-specific cytotoxicity could be attributed to enterotoxin gene presence. A multi-way ANOVA with interaction effects revealed significant associations between cytotoxicity and the presence of the *hbl* operon (p = 0.025), *cytK-2* (p = 1.7e-11), and *panC* group (p = 1.3e-10). However, no significant interaction was found between enterotoxin genes and *panC* group (p = 0.34). Detailed statistical analyses results are available in the Supplemental Material Table S2.

Post-hoc Tukey’s HSD analyses indicated that isolates from *panC* groups I, IV, and V were significantly more cytotoxic than those from other groups. Specifically, group I isolates exhibited significantly higher cytotoxicity compared to groups II, III, and VI, and group IV and V isolates showed significantly higher cytotoxicity than groups II and VI. Detailed p-values for pairwise comparisons are provided in Supplemental Material Table S3. These findings are consistent with previous studies by Guinebretière et al. 2010 and Miller et al. 2018, though group I isolates were not included in the former study (10, 11).

Group IV includes *B. cereus sensu stricto* (*s.s.*), a known foodborne pathogen, as well as commercially used *B. thuringiensis* strains that produce insecticidal crystal proteins encoded by *cry* genes. The pathogenic potential of *B. thuringiensis* strains and their use in agriculture has been contested (39–42). Our study analyzed 43 isolates harboring *cry* genes, indicating that these isolates belong to biovar Thuringiensis. These isolates had a mean cytotoxicity of 0.79 ± 0.36, higher than the overall mean for *B. cereus s.l.* (0.51 ± 0.35) and comparable to group IV isolates (0.76 ± 0.36) (Table 1). This emphasizes the importance of safety evaluations for individual *B. thuringiensis* strains intended for commercial use.

Interestingly, group VII isolates did not exhibit uniformly high cytotoxicity despite *B. cytotoxicus* being linked to foodborne outbreaks (43) and the greater cytotoxic potential of CytK-1 compared to CytK-2 (19). Rather, group VII isolates displayed a bimodal cytotoxicity distribution, with half showing cytotoxicity around 0.7 and the other half exhibiting cytotoxicity around 0 (Fig. 3). These results align with earlier studies that reported variability in group VII isolate cytotoxicity against Vero cells (30, 44–46). Further analysis of additional group VII isolates could provide more clarity on the safety of *B. cytotoxicus*.

Residual variation in the ANOVA could be explained by differences in the enterotoxin protein sequences that enhance toxin efficacy, variations in overall enterotoxin expression levels, or contributions from other virulence factors, such as phospholipases like sphingomyelinase (47), and phosphatidylinositol-specific phospholipase C (PI-PLC) (48, 49), metalloproteases like InhA1 and NprA (24, 50), and hemolysins like hemolysin I (cereolysin O), hemolysin II, and hemolysin III (51–53). Further studies on these factors may provide additional insights into cytotoxicity variation within *B. cereus s.l*.

### Enterotoxin gene presence is a sensitive but nonspecific predictor of cytotoxicity

Our findings and prior studies demonstrate that *B. cereus s.l.* strains can harbor enterotoxin genes without exhibiting cytotoxicity (11, 27, 29, 41). This suggests that the presence of enterotoxin genes alone may not reliably predict cytotoxicity. To evaluate this, we applied logistic regression to test whether cytotoxicity could be accurately predicted by the presence of specific enterotoxin genes (*hbl*, *nhe*, *cytK-1*, and *cytK-2*). Reliable predictors would exhibit high sensitivity (correctly identifying cytotoxic isolates) and high specificity (correctly identifying non-cytotoxic isolates). We calculated the accuracy, precision, sensitivity, and specificity for each enterotoxin gene, with high-quality predictors having values close to 1.

All individual *nhe* and *hbl* genes, as well as the entire *nheABC* and *hblACD* operons, had sensitivity values near 1 and specificity values near 0 (Fig. 4A). This indicates that highly cytotoxic isolates were almost always correctly classified as possessing the gene. However, many non-cytotoxic or low-cytotoxicity isolates were misclassified, resulting in frequent type I errors. Among the enterotoxin genes, *cytK* genes demonstrated the highest specificity values, near 1 (Fig. 4A). *cytK-2* also showed the largest difference in cytotoxicity between isolates with and without the gene (Fig. 4A).

**FIG 4.**
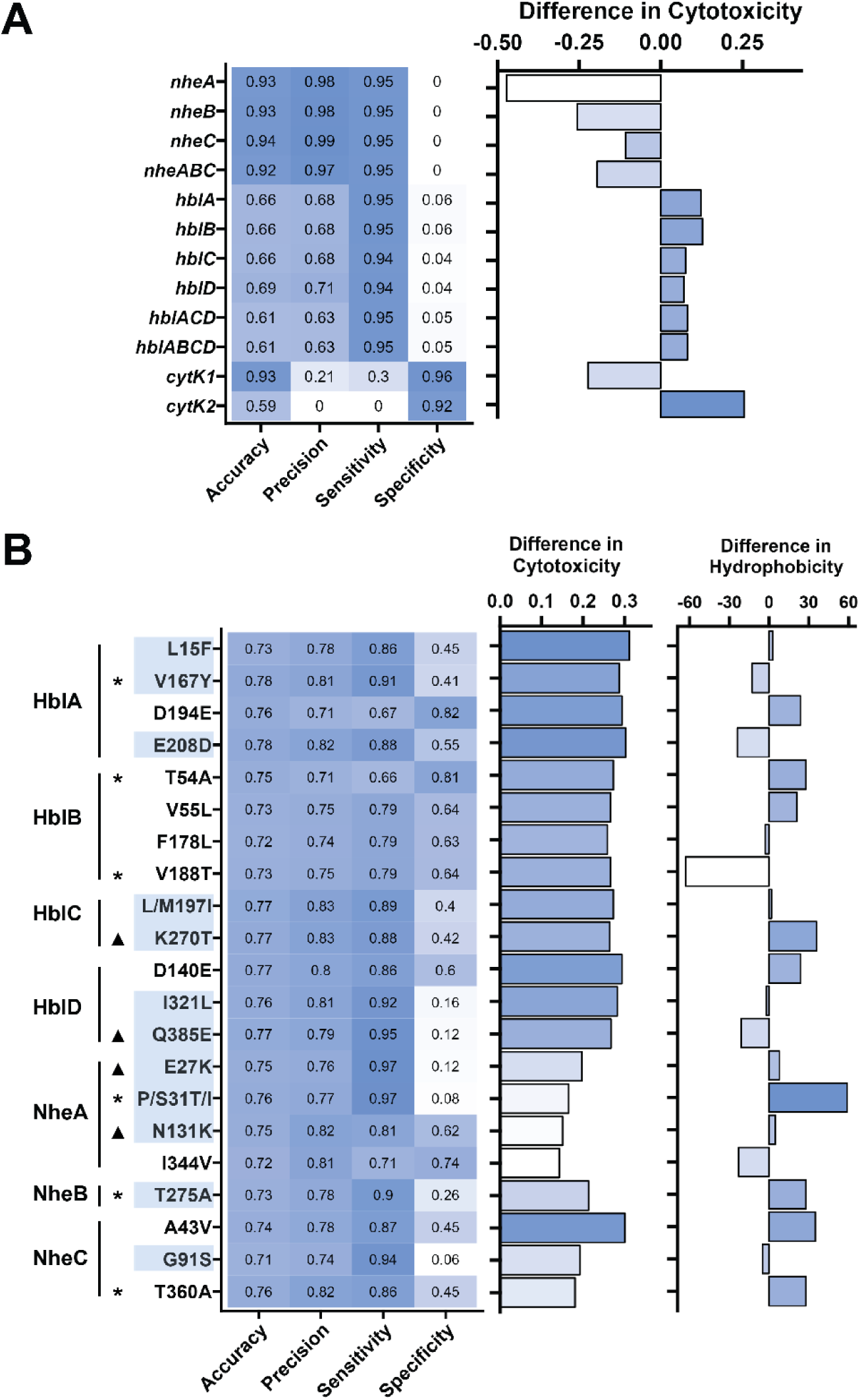
Potential of enterotoxin genes and SNPs for prediction of cytotoxicity. **A.** Statistics for prediction of cytotoxicity based on enterotoxin gene presence, and the mean difference in cytotoxicity between isolates with and without the gene. Negative cytotoxicity values indicate that isolates with the gene were less cytotoxic than isolates without the gene. **B.** Statistics for prediction of cytotoxicity based on the SNPs detected in enterotoxin genes. The mean difference in cytotoxicity between isolates with either SNP variant and the difference in hydrophobicity between SNP amino acids are shown. The hydrophobicity index was derived from Monera et al. 1995 (56), where the scale was normalized to glycine (0) and the most hydrophobic residue (100). SNP are annotated as follows: amino acid associated with low cytotoxicity, position in the protein, amino acid associated with high cytotoxicity. SNPs are grouped by their respective enterotoxins. SNPs associated with cytotoxicity, but not phylogenetic group, are highlighted in light blue. For SNPs L/M197/I and P/S31T/I, where multiple amino acid substitutions were detected, the difference in hydrophobicity is shown for the first listed substitutions (L197I and P31T). Triangles next to SNP labels denote variants in which amino acid charge is different between the two variants. Asterisks next to SNP labels denote variants in which amino acid hydrogen bonding ability is different between the two variants.

Isolates with *nhe* and *cytK-1* genes exhibited lower cytotoxicity than isolates lacking these genes (Fig. 4A). This could be a result of small sample sizes, as only 7 genomes lacked *nhe* and 9 contained *cytK-1* (Table 1). The small sample size also likely inflates accuracy and precision values for these genes.

Enterotoxin gene presence is an imperfect predictor of cytotoxicity due to variability in gene expression and function. *Nhe* genes, for instance, are nearly ubiquitous in *B. cereus s.l.* (10, 12, 35), but are not constitutively expressed (11, 27, 29, 41). Although Nhe proteins are commonly detected in supernatant, not all isolates producing Nhe are cytotoxic (11, 29). Conversely, in some strains, Nhe proteins alone are sufficient to cause cytotoxicity and illness (11, 54). These inconsistencies could be explained by genetic variations in enterotoxin sequences that may alter their expression or functionality, or by *in vitro* assays not fully capturing the cytotoxic potential of diverse isolates due to technical limitations or differences in experimental conditions.

### SNPs in enterotoxin gene sequences are sensitive and specific predictors of cytotoxicity

Existing markers for predicting *B. cereus s.l.* cytotoxicity rely on the presence of entire genes or combinations of genes (55). For instance, a recent study has proposed seven pathogenicity biomarker genes, including transcription factors and understudied genes, as differentially expressed between clinical and environmental isolates (55). However, single nucleotide polymorphisms (SNPs) within enterotoxin genes have not been explored as potential predictors of cytotoxicity.

To identify higher-quality predictors of cytotoxicity than gene presence alone, we analyzed nonsynonymous SNPs in enterotoxin genes that alter protein sequences and potentially impact function. Using BTyper3, we extracted *hbl*, *nhe*, *cytK-1,* and *cytK-2* sequences, which were then aligned. Logistic regression was employed to identify SNPs that met the following criteria: (i) significantly correlated with cytotoxicity after Bonferroni correction, (ii) predicted cytotoxicity with accuracy and precision greater than 70%, and (iii) caused a nonsynonymous amino acid substitution. Phylogenetic group data was incorporated in the logistic regression to assess whether identified SNPs were associated with cytotoxicity independently of phylogeny. SNPs were annotated in the format: amino acid variant associated with lower cytotoxicity, followed by the position in the amino acid alignment, followed by the amino acid variant associated with higher cytotoxicity.

A total of 21 nonsynonymous SNPs significantly associated with cytotoxicity were identified (Supplemental Table S4). Of these, 13 were found in *hbl* genes and 8 in *nhe* genes (Fig. 4B). No SNPs were identified in *cytK-1* or *cytK-2*, likely due to low sequence variation in these genes (Supplemental Material Fig. S1). Twelve of 21 SNPs were significantly correlated with cytotoxicity, but not phylogenetic group, indicating their potential utility as independent markers of cytotoxicity (Fig. 4B, Table S4). Fifteen of the 21 exhibited higher specificity for predicting cytotoxicity compared to the presence of *nhe* and *hbl* genes (Fig. 4). The mean difference in normalized cytotoxicity between strains with or without a SNP variant was 0.246 ± 0.0540. These findings demonstrate that nonsynonymous SNPs in *hbl* and *nhe* genes are sensitive and specific predictors of cytotoxicity, surpassing gene presence alone in predictive power.

### SNPs in enterotoxin genes, *panC* group, and *hbl* gene presence are the key predictors of cytotoxicity in a random forest regression

To complement our logistic regression results, we performed random forest regression analyses to identify the most important predictors of cytotoxicity using phylogenetic group, enterotoxin genes, and SNPs. Predictor importance was assessed using the permutational variable importance (PVI), calculated by randomly shuffling the data and measuring its impact on the model error. Higher PVI values indicated greater predictor importance.

SNPs and *panC* group had the three highest mean PVI values. The SNP NheC A43V was the top-ranked variable, demonstrating that its inclusion improved model performance more than any other predictor (Fig. 5). *panC* group was the second most important variable, consistent with its significant correlation with cytotoxicity (Fig. 3).

**FIG 5.**
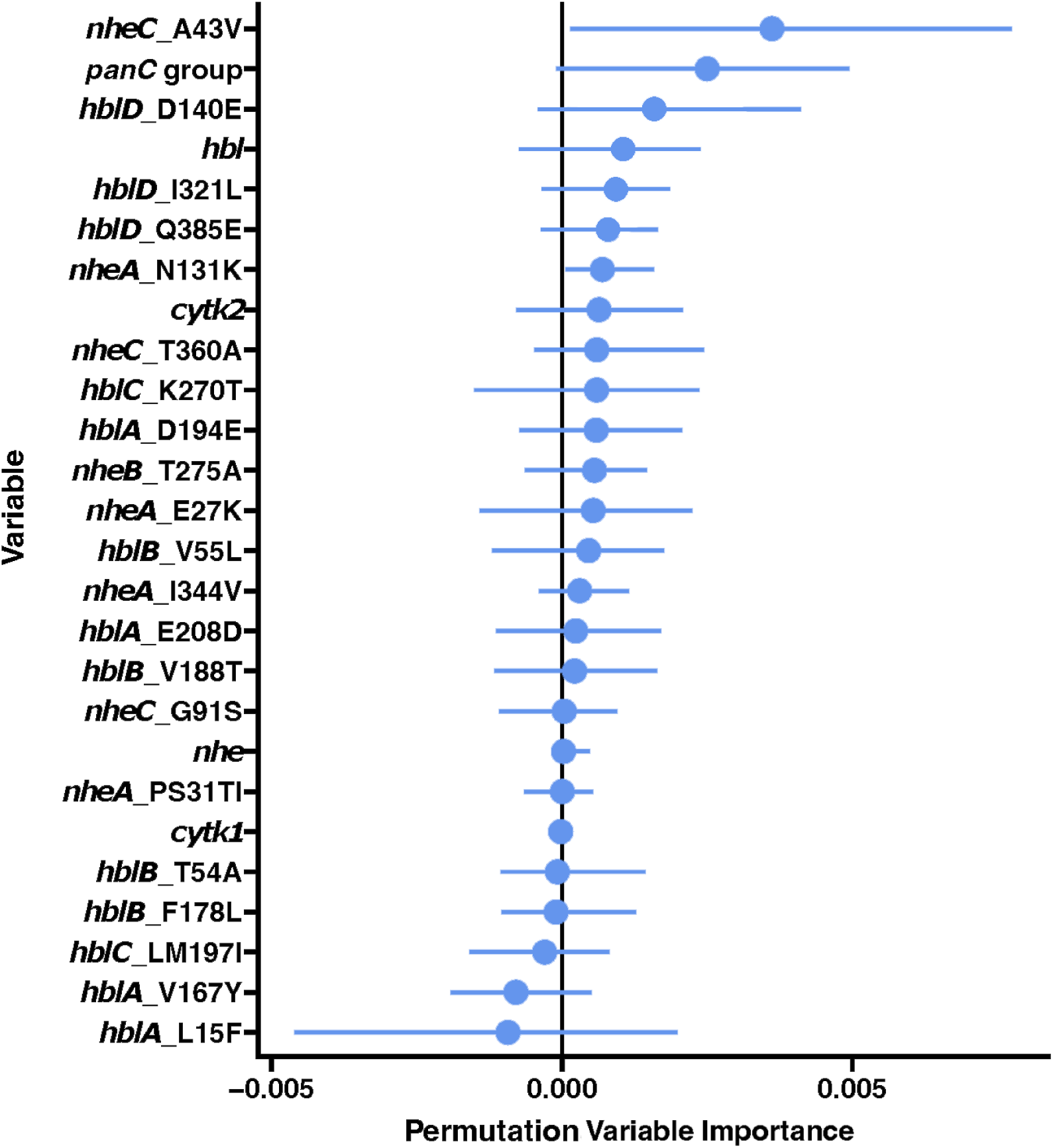
Permutation variable importance identified through random forest regression analyses. Circles represent the mean permutation variable importance of phylogenetic group, enterotoxin genes, and SNPs in enterotoxin gene sequences for prediction of cytotoxicity, based on 100 calculation repetitions. Lines signify a 95% confidence interval.

The *hbl* operon ranked 4^th^ out of 26 variables, and *cytK-2* ranked 8th (Fig. 5). These findings align with our multi-way ANOVA and logistic regression results, which showed *hblACD* and *cytK-2* to be significantly correlated with cytotoxicity and *cytK-2* to have higher specificity than *nheABC* (Fig. 4). The *nhe* operon and *cytK-1* had mean PVI values near zero, suggesting their inclusion did not improve the model performance. This was expected for the *nhe* operon, as it is present in nearly all isolates resulting in minimal variability (Table 1).

Some SNPs had slightly negative PVI values, indicating they were less predictive of cytotoxicity than random data. This may result from splitting importance values among correlated variables. For instance, SNPs like HblA L15F, which are associated with both cytotoxicity and *panC* group, likely had deflated values due to the inclusion of *panC* group in the model.

### Identified SNPs may alter protein structure and function

Nonsynonymous amino acid mutations can alter protein function by reshaping the protein structure or modifying binding residues. Nhe and Hbl proteins are rich in long alpha helices connected by linkers (Fig. 6 and 7). In total, 14 of 21 SNPs occur within alpha helices, one SNP occurs in the signal peptide region, and 6 SNPs occur within linker regions (Fig. 6 and 7). Nhe proteins have 4 SNPs within an alpha helix and 4 SNPs within linker regions. Conversely, Hbl proteins have 11 SNPs within alpha helices, one SNP within the signal peptide, and only one SNP within a linker region. Our identified SNPs could affect the protein secondary or tertiary structure in these regions and affect protein function and cytotoxicity downstream.

**FIG 6.**
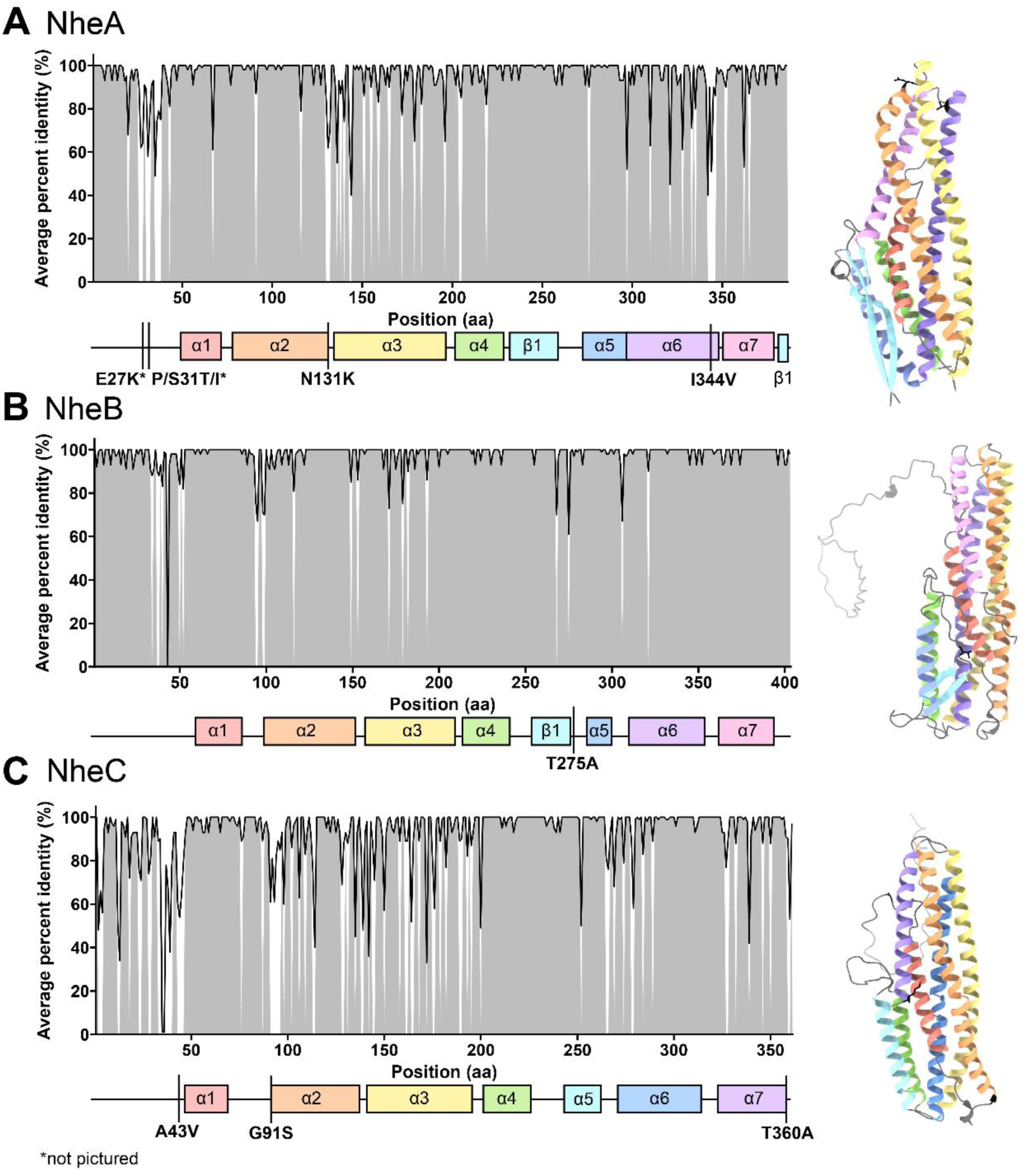
Nhe protein sequence homologies and structures with highlighted SNPs. Amino acid percent identity values across the protein sequence are plotted above linear protein maps. Gray areas in percent identity plots indicate regions with >90% identity across sequences. Vertical lines in protein maps indicate the location of SNPs associated with cytotoxicity. Protein secondary structures are colored in rainbow order from N to C-terminus, as on the linear map. SNPs are indicated on the structure in black. Signal peptides are absent in structure A, and shown in light gray in structures B and C.

**FIG 7.**
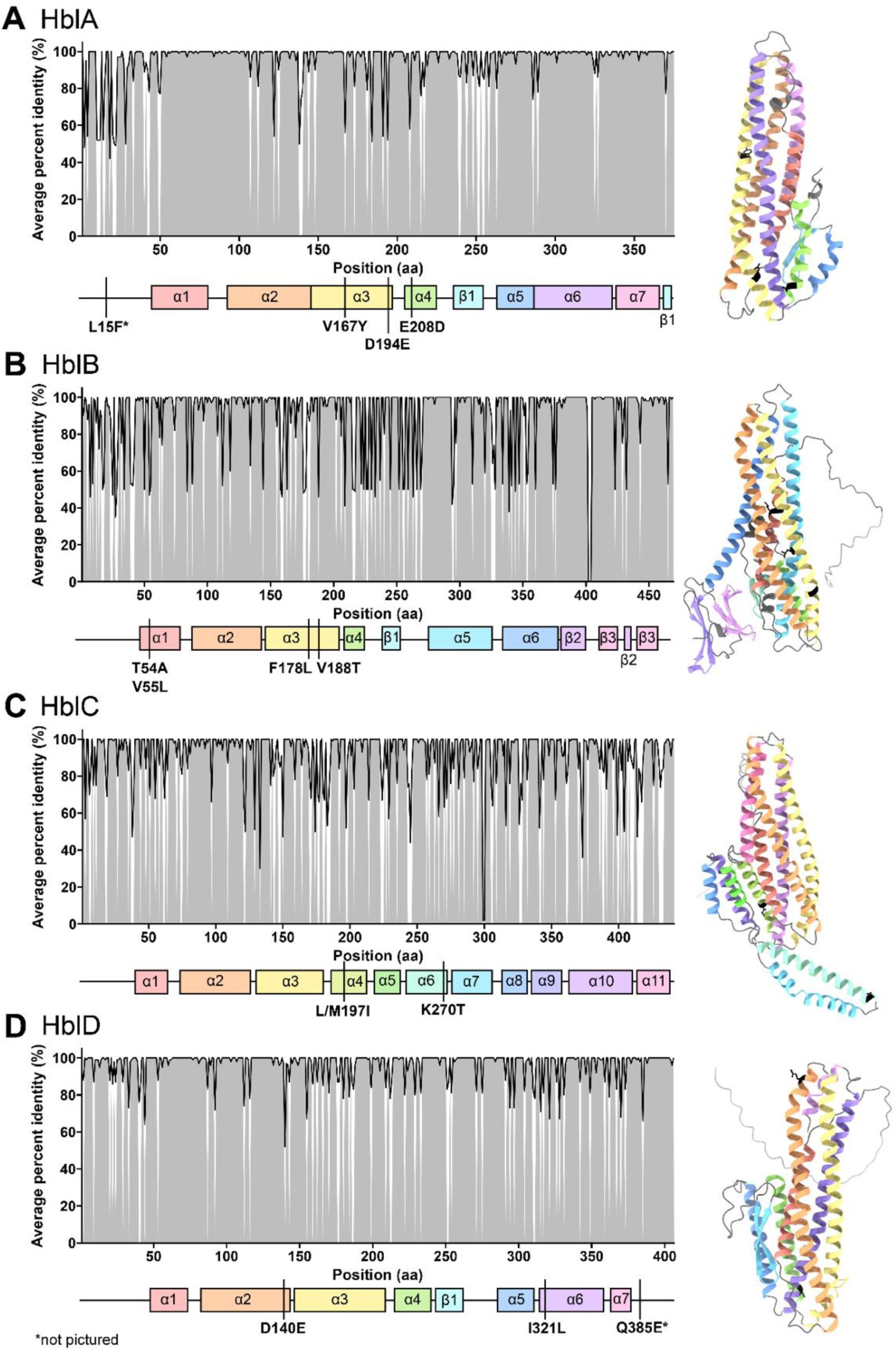
Hbl protein sequence homology and structure with highlighted SNPs. Amino acid percent identity values across the protein sequence are plotted above linear protein maps. Gray areas in percent identity plots indicate regions with >90% identity across sequences. Vertical lines in protein maps indicate the location of SNPs associated with cytotoxicity. Protein secondary structures are colored in rainbow order from N to C-terminus, as on the linear map. SNPs are indicated on the structure in black. Signal peptides are absent in structure A, and shown in light gray in structures B, C, and D.

Amino acid hydrophobicity strongly influences protein structure and function. Therefore differences in hydrophobicity can be used as a proxy for the magnitude of protein alteration between SNPs. For our data, we calculated the difference in hydrophobicity between the two residues by subtracting the hydrophobicity of the residue associated with lower cytotoxicity from the residue associated with higher cytotoxicity, by using the hydrophobicity index presented by Monera et al. 1995 (56). The scale was normalized such that glycine was classified as neither hydrophobic nor hydrophilic and had a value of 0, and the most hydrophobic residue had a value of 100. A negative difference in hydrophobicity indicates that the amino acid associated with lower cytotoxicity is more hydrophobic than the amino acid associated with higher cytotoxicity. The average absolute difference in hydrophobicity between amino acids associated with elevated cytotoxicity and those associated with lower cytotoxicity is 29.9 ± 32.4. SNPs with extraordinarily large differences in hydrophobicity between the two residues included HblB V188T (-63) and NheA P31T, P31I and S31I (59, 145, and 104, respectively). Large differences in hydrophobicity indicate that the amino acid substitution likely strongly affects the enterotoxin structure and therefore function. For example, a more hydrophobic residue may increase the strength of protein-protein interactions between enterotoxin subunits, as many protein-protein interfaces include hydrophobic residues (57). Specifically as a member of the α-helical pore-forming toxin family, the Nhe complex requires a hydrophobic region in NheC, likely for membrane insertion, to cause cytotoxicity against Vero cells (13, 16). Therefore, changes to the hydrophobicity of residues in this region may lead to an inability of NheC to insert into the membrane. Though the Hbl complex resembles that of Nhe, the analogous hydrophobic region on HblA is unnecessary to cause cytotoxicity (16), so it remains unknown which regions of Hbl are required for membrane insertion.

We also examined other major changes of the SNPs, including differences in charge or hydrogen bonding ability. These alterations could cause substantial changes in protein structure and bonding abilities with host receptors and membranes or other enterotoxin subunits. Changes in charge are exhibited for 4 SNPs: HblC K270T, HblD Q385E, NheA E27K, and NheA N131K (Fig. 4). Changes in hydrogen bonding ability are exhibited for 7 SNPs: HblA V167Y, HblB T54A, HblB V188T, NheA P31T and S31I, NheB T275A, NheC G91S, and NheC T360A (Fig. 4).

Though we only investigate enterotoxins and their variants in this paper, alternative markers of cytotoxicity could include the expression of enterotoxins and variants of other virulence factors. As discussed by previous studies, enterotoxin expression levels add an additional layer of complexity to understanding *B. cereus s.l.* virulence (27, 58, 59). Expression is controlled by several transcription factors that respond to environmental signals associated with the intestine (58–60). It is also possible that the SNPs identified here (as well as synonymous SNPs) could alter mRNA secondary structure and have downstream effects on translation and expression.

### Identified SNPs are located within key virulence-related regions of enterotoxins

The identified SNPs are positioned within protein regions critical for enterotoxin function, potentially affecting their transport, host-cell targeting, and pore-forming activity. Signal peptides guide enterotoxins to the host cell membrane via the Sec secretion pathway (61). Based on the signal peptide region predicted by SignalP in the UniProt Automatic Annotation pipeline, HblA L15K is located within the signal peptide region, and NheA E27K resides at the predicted signal peptide cleavage site. Both SNPs involve charge-altering substitutions that could influence secretion efficiency or processing.

The Nhe proteins interact in a sequential manner to form a cytotoxic pore complex on the host membrane. NheB and NheC bind the host membrane first and form a pro-pore complex, facilitated by transmembrane regions (62, 63). One of the identified SNPs, NheB T275A, lies within a predicted transmembrane region, and therefore may affect insertion (64, 65). Once NheB and NheC are bound, NheA interacts with the NheB-C complex and undergoes a conformational change to form the final transmembrane pore (62, 64, 65). SNPs in flexible loop regions may influence this process. For example, NheA N131K resides in a region important for the transition to the full pore conformation, potentially affecting the rigidity or structural changes required for pore formation (65).

The Hbl proteins operate in a similar pore-forming manner, but their specific mechanisms are less well understood. HblA is the host-binding protein, and HblD and HblC are the lytic components. HblA binds to the host membrane, after which HblD can complexes with HblA, either in solution or at the membrane (15, 66). Finally, HblC joins the HblA-D complex, likely inducing a conformational change to complete pore formation (15). Although a transmembrane region has been predicted in HblA, none of the identified SNPs are located within this region (63). Additionally, some *hbl* operons include a fourth gene, *hblB,* which encodes a protein homologous to HblA, but with a distinct regulatory role. HblB cannot substitute for other Hbl proteins inhibits cytotoxicity by outcompeting HblA for binding to HblD (67). Four SNPs were identified in *hblB*.

Despite the structural and functional insights into Nhe proteins, less is known about the precise residues critical for Hbl protein function. Few studies have clarified the exact mechanisms of Hbl activity or identified specific binding residues, making it challenging to contextualize the potential impacts of the identified SNPs in Hbl proteins on cytotoxicity (14, 15, 66, 68). This gap highlights the need for further research into the molecular interactions governing Hbl-mediated virulence.

### Conclusions and study limitations

Our study demonstrates that nonsynonymous SNPs in enterotoxin gene sequences serve as more effective predictors of cytotoxicity than enterotoxin gene presence or *panC* phylogenetic group alone. This finding highlights the potential of SNP-based markers to enhance the identification of *B. cereus s.l.* isolates with elevated risk for toxicoinfection. Notably, SNPs such as NheC A43V, NheB T275A, and HblA L15K are located in critical for protein function, such as signal peptides, transmembrane domains, and regions involved in pore formation. These SNPs likely influence key processes such as secretion efficiency, membrane insertion, and conformational changes required for cytotoxic activity. The ability of SNPs to predict cytotoxicity with greater specificity and sensitivity than gene presence highlights their promise as robust biomarkers for strain-based risk assessment in food safety.

However, our study utilizes an *in silico* approach, which does not capture all factors contributing to toxicoinfection. For example, the ability of *B. cereus s.l.* isolates to adapt to the human gastrointestinal tract’s microenvironments, interact with the resident microbiota, or evade the human immune response likely influence cytotoxicity but were not assessed in this study. Furthermore, the precise functional impacts of the identified SNPs on protein activity remain to be experimentally validated to confirm the mechanistic role of these SNPs in virulence.

## MATERIALS AND METHODS

### Isolates used in the study

A total of 270 *B. cereus s.l.* isolates were included in this study to represent a variety of isolation sources and all eight *panC* phylogenetic groups. The list of isolates with corresponding metadata is available in Supplemental Material Table S1. All isolates were stored at -80 °C in brain heart infusion broth (BHI, BD Difco) supplemented with 20% (vol/vol) glycerol. For each experiment, BHI broths were inoculated with a single colony. Inoculated BHI broths were incubated at 37°C for 9-12 hours without shaking. Cultures were washed in PBS and adjusted to approximately 1 x 10^6^ CFU/ml based on a *B. cereus* ATCC 14579 OD-CFU standard curve.

### Determining early stationary growth phase

Adjusted cultures were deposited into wells of 96-well plates in three technical replicates. Inoculated plates were incubated at 37°C without shaking. Optical density (OD_600_) was measured after a 15 second orbital shake every 30 minutes for 24 hours using a microplate reader (Synergy Neo2, Biotek).

OD_600_ data of three replicates per isolate were averaged and a logistic growth model was fitted using the R package “growthcurver” v0.3.0 (69). To ensure an accurate fit, the death phase data were removed (defined as any data points that deviated by >0.2 units from the growth maximum). Early stationary phase was defined as the intercept of the tangent lines for exponential phase and stationary phase + 2 hours (36). The times needed for each isolate to reach early stationary phase were used as the isolate incubation times for supernatant harvesting to ensure that cell density and quorum-sensing-regulated enterotoxin expression reached its peak (36, 70, 71).

### Supernatant harvesting and filtering

Adjusted cultures were incubated without shaking at 37°C until each isolate reached early stationary phase. Cultures were centrifuged (Avanti Centrifuge J-26 XPI) at 16,000 g for 2 minutes and supernatants were filtered using 0.2 µm cellulose acetate filters (VWR) and stored at -80°C until use in cytotoxicity experiments.

### Cytotoxicity in a Caco-2 cell model

Caco-2 cells were purchased from ATCC and cryopreserved in Eagle’s Minimum Essential Media (Corning) with 20% fetal bovine serum (FBS, VWR Life Science Seradigm), supplemented with 5% DMSO, in liquid nitrogen. Caco-2 cells were grown at 37°C, 95% humidity, and 5% CO_2_ for 4-6 days. Cells were incubated in completed EMEM supplemented with 20% FBS, and 1% penicillin-streptomycin solution (Gibco), which was changed every 2 days. Cells were detached for splitting using 0.25% trypsin-EDTA (Gibco). Before experiments proceeded, cell cultures were tested for mycoplasma contamination using the Venor GeM Mycoplasma Detection Kit (Sigma Aldrich).

Caco-2 cells were seeded at 1 x 10^4^ cells per well in 96-well plates for assessment of cell-free supernatant cytotoxicity, which was carried out using WST-1 cell proliferation assays (37). Plates were incubated for 3-4 days at 37°C, 95% humidity, and 5% CO_2_ to allow for cell attachment and growth to approximately 100% confluency. Medium was changed the day before a cytotoxicity experiment to ensure a high baseline cellular metabolism. Wells were treated with 15% (vol/vol) of either test bacterial cell-free supernatant, *B. cereus* ATCC 14579 supernatant (positive cytotoxicity control), or sterile BHI (negative cytotoxicity control). Each isolate supernatant was tested in six technical replicates per plate in two biological replicates (i.e., using supernatants collected in two independent experiments). After treatment, plates were incubated for 15 minutes to allow for intoxication. Then, 10% (vol/vol) of WST-1 dye was added to each well, and cells were incubated for an additional 105 minutes to allow for metabolic conversion of WST-1 dye. The intensity of the metabolized WST-1 dye corresponded with the level of metabolic activity. The color intensity was measured based on the dye absorbance at 450 nm and 690 nm.

Absorbances were calculated by subtracting the background signal measured at 690 nm from the signal measured at 450 nm. Absorbances for test wells were then normalized to the controls of their individual plates using min-max normalization (38) with the following formula.

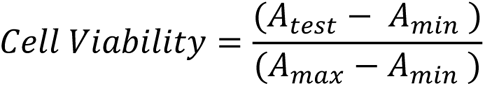

A_test_ represents the average experimental absorbance of the six technical replicates for a single isolate. A_min_ represents the average absorbance of the BHI negative control replicates. A_max_ represents the average absorbance of the six *B. cereus* ATCC 14579 positive control replicates. The assay was performed in biological duplicate using supernatants from two distinct bacterial colonies and two separate Caco-2 plates. The resulting values were averaged to obtain the final normalized cytotoxicity value.

### Whole-genome sequencing and sequence analyses

A total of 240 isolates that had not been previously whole-genome sequenced (3, 11) were whole-genome sequenced in this study. Isolates were grown in BHI broth at 37°C for 6 hours. Genomic DNA was extracted from grown culture suspensions using the E.Z.N.A. Bacterial DNA Kit (Omega), followed by the glass bead-beating protocol provided by the kit manufacturer. The final DNA concentration was adjusted to 50 ng/µl using nuclease-free water. Extracted DNA was used for Nextera XT library preparation and Illumina sequencing using with 150 bp paired-end reads using the NextSeq platform (Wright Labs, PA).

The quality of sequencing reads was examined with FastQC v0.11.5 (72, 73), using the default settings. Adapters and reads with quality scores below 3 or lengths shorter than 36 bp were removed using Trimmomatic v0.36 with leading and trailing options enabled. Trimmed reads were then assembled via *de novo* assembly with SPAdes v3.13.1 (74) with k-mer lengths 99 and 127 bp, and the --isolate option. The assembly quality was checked using the default settings of QUAST v4.6.1 (75), and average coverage of each genome was determined using BWA v0.7.17 (76) and Samtools v1.7 (77). Core genome SNPs were identified with kSNP v2.13 using a k-mer length of 21 bp (78) and used to construct a maximum likelihood phylogenetic tree with IQ-TREE 2 v2.2.0 (79), using the generalized time-reversible model with a gamma distribution and ascertainment bias correction (GTR+G+ASC) and 1000 ultrafast bootstraps (80–82). The phylogenetic tree was visualized using ggtree v3.6.2 (83). Assembled draft genomes were analyzed using BTyper v3.2.0 (33) to identify taxonomic species using an Average Nucleotide Identity (ANI) method, detect virulence genes, and extract *nhe*, *hbl*, *cytK-1,* and *cytK-2* gene sequences using 70% coverage and 50% amino acid identity cut-offs to ensure inclusive detection of genes that were present but disrupted by assembly gaps.

### Identification of genetic markers predictive of cytotoxicity

All analyses were done using Python v3.8 and “statsmodel” v0.12.2 (84) unless otherwise stated. To determine if cytotoxicity could predict enterotoxin gene presence, a logistic regression model with the maximum likelihood estimation method was fitted to normalized cytotoxicity data and enterotoxin gene presence. A confusion matrix, accuracy, and precision score were calculated using the “scikit-learn” v0.24.1 Python module (85). Sensitivity was calculated by dividing the number of true positives by the sum of true positives and false negatives. Specificity was calculated by dividing the number of true negatives by the sum of true negatives and false positives.

To identify SNPs predictive of cytotoxicity, enterotoxin gene sequences were aligned using MUSCLE v3.8.425 (86). A logistic regression model as described above was fitted to cytotoxicity and *panC* group data (considered covariates) and nucleotide presence at each position in the alignment. A nucleotide was defined as “present” if the nucleotide of interest was in the position being examined. A nucleotide was defined as “absent” if a different nucleotide was found in this position. Isolates with a gap in the position were removed from the analysis of that position. Only positions with between 20-80% conservation were included in the analyses to identify common, but not hyperconserved nucleotides. These limits were used to reduce misclassifications. Confusion matrices, accuracy, precision, sensitivity, and specificity scores were calculated as previously described. Two-tailed p-values for the t-statistics of each covariate in explaining SNP presence or absence were calculated. Bonferroni correction was applied to the p-values to correct for multiple comparisons. SNPs where all isolates were classified as only positive or only negative classes, SNPs with lower than 70% accuracy or precision, and SNPs that did not result in a nonsynonymous amino acid substitution were filtered out.

In addition to logistic regression, a random forest regression model was developed, incorporating *panC* group, virulence genes, and SNPs to predict cytotoxicity. The “mlr3” package v3.16.0 (87) was employed for model training and evaluation. A ten-fold cross-validation was utilized to tune two hyperparameters (number of trees (ntree) and number of variables for each split (mtry)) by minimizing mean squared error of the model (88). Permutational variable importance was computed 100 times and averaged to identify informative variables for the regression (89).

### Additional statistical analyses

Statistics were calculated using R “stats” v4.2.2. The Shapiro-Wilk normality test was used to determine whether cytotoxicity values were normally distributed. A multi-way ANOVA with an interaction term was performed for enterotoxin genes (*nhe* and *hbl* operons, *cytK-1*, and *cytK-2*) and *panC* group to test significant association with cytotoxicity. A Tukey’s honest significant differences post-hoc test was used to compare significant differences between the average cytotoxicity values of each *panC* group. Additional test statistics are reported in Table S2.

### Protein structure accession numbers

Crystal structures were downloaded from PDB for NheA (4K1P) and HblA (2NRJ). AlphaFold structures were downloaded from UniProt for NheB (Q09KI4_BACCE), NheC (Q09KH9_BACCE), HblB (Q9REG5_BACCE), HblC (O05491_BACCE), and HblD (O05492_BACCE).

### Data availability statement

Whole-genome sequences used in this study are available through NCBI, and their accession numbers are listed in Supplementa. Scripts for analyses can be accessed at https://github.com/cassprince/B-cereus-cytotoxicity-SNPs. Raw cytotoxicity data will be made available upon request.

## ACKNOWLEDGMENTS

This work was supported by the United States Department of Agriculture (USDA) National Institute of Food and Agriculture (NIFA) project 2019-67017-29591, Hatch Appropriations under Project PEN04853 and Accession 7005519, and the Multistate Project 4666.

## SUPPLEMENTAL FIGURE

**FIGURE S1** CytK protein sequence homology. Amino acid percent identity values across the protein sequence are plotted above linear protein maps. Gray areas in percent identity plots indicate regions with >90% identity across sequences.

## SUPPLEMENTAL TABLES

**TABLE S1** Isolates used in the study.

**TABLE S2** Multi-way ANOVA with interaction effects Tukey HSD p-values (adjusted for multiple comparisons) for testing the association of cytotoxicity and enterotoxin gene presence.

**TABLE S3** Multi-way ANOVA with interaction effects Tukey HSD p-values for comparing the cytotoxicity of *panC* phylogenetic groups.

**TABLE S4** Bonferroni corrected p-values from the logistic regression associating SNPs with cytotoxicity and *panC* phylogenetic group.

